# Assessing variant effect predictors and disease mechanisms in intrinsically disordered proteins

**DOI:** 10.1101/2025.04.01.646619

**Authors:** Mohamed Fawzy, Joseph A. Marsh

## Abstract

Intrinsically disordered protein regions (IDPRs) are central to diverse cellular processes but present unique challenges for interpreting genetic variants implicated in human disease. Unlike structured protein domains, IDPRs lack stable three-dimensional conformations and are often involved in regulation through transient interactions and post-translational modifications. These features can affect both the distribution of pathogenic variants and the performance of computational tools used to predict their effects. Here, we systematically assessed the distribution of pathogenic *vs* benign missense variants across disordered, intermediate, and structured protein regions in the human proteome. Pathogenic variants were notably depleted in IDPRs yet were associated with distinct molecular mechanisms, particularly dominant gain- and loss-of-function effects. We evaluated 33 variant effect predictors (VEPs), revealing widespread reductions in sensitivity for pathogenic variants in IDPRs—despite high AUROC scores largely driven by accurate benign variant predictions. We also observed substantial discordance among VEP classifications in disordered regions, underscoring the need for region-aware thresholds and disorder-informed prediction strategies. Incorporating features reflective of IDPR biology, such as transient interaction motifs and modification sites, may enhance the accuracy and interpretability of future tools.

## Introduction

Intrinsically disordered protein regions (IDPRs) lack stable secondary or tertiary structure under physiological conditions, instead adopting flexible conformations that can shift upon binding to specific partners or in response to cellular cues [1–3]. This structural plasticity enables IDPRs to act as molecular hubs in diverse cellular processes, including transcriptional regulation, signal transduction, and protein–protein interactions [4]. Their binding is typically characterised by high specificity but low affinity, allowing for transient, reversible interactions [2,5], while their conformational flexibility facilitates extensive post-translational modification (PTM), fine-tuning function in response to environmental changes [6–9]. IDPRs are especially prevalent in eukaryotic proteins, with around 30–40% containing long disordered regions (>30 residues) [10], and their sequence composition—enriched in polar and charged residues, and depleted in hydrophobic residues—prevents stable folding, favouring a dynamic equilibrium of conformational states [1,11,12]. These properties make IDPRs central to regulatory networks and sensitive to perturbations, which may explain their involvement in a wide range of human diseases, including cancer, cardiovascular, neurodegenerative, and prion disorders [9].

Variant effect predictors (VEPs) are computational tools designed to estimate the potential impact of genetic variants, particularly on human health and disease. There are many different VEPs, each using distinct types of information—such as evolutionary conservation, structural features, allele frequencies, or functional assay data—and varying widely in their algorithms, training data, and outputs [13]. While VEPs are widely used, relatively little attention has been given to how their performance varies depending on whether a variant is in a structured or disordered region. Despite the functional importance of IDPRs, these regions might pose significant challenges for VEPs. Unlike structured protein domains, IDPRs tend to be less evolutionarily conserved and lack the fixed secondary and tertiary structures that some VEPs rely on as predictive features. As a result, VEPs may struggle to accurately classify pathogenic variants in these regions. Previous studies have shown that standard predictors such as SIFT and PolyPhen-2 exhibit a reduction of more than 10% in sensitivity for pathogenic variants in IDPRs compared to structured regions [14]. However, a systematic evaluation of VEP performance across different protein structural contexts remains largely unexplored.

Previously, we showed that IDPRs significantly influence the apparent performance of missense VEPs, often leading to inflated area under the receiver operating characteristic curve (AUROC) values [15]. This elevation in AUROC stems from the fact that disordered regions are enriched with putatively benign missense variants, which are typically under weaker evolutionary constraint and thus easier for VEPs to correctly classify as benign. However, this can be misleading: while it is not technically incorrect, it primarily reflects the ease with which VEPs handle these straightforward benign classifications rather than their proficiency in detecting disease-causing mutations. This distinction is critical, as it suggests that the impressive AUROC values observed in proteins with large disordered regions may not indicate robust performance in identifying pathogenic variants, which is often the primary concern in clinical and research settings.

Building on this observation, here we have further investigated the occurrence of pathogenic missense variants and the predictive performance of VEPs in IDPRs. First, we systematically classified residues across the human proteome to assess the distribution of pathogenic and benign variants in disordered versus structured regions. We evaluated 33 VEPs, spanning clinical-trained, population-tuned, and population-free categories, to determine their effectiveness in identifying pathogenic variants within IDPRs. Our analysis reveals significant differences in variant distribution and VEP performance across structural contexts, highlighting the limitations of current predictors in capturing the subtle functional impacts of variants in disordered regions. Furthermore, we explored the molecular mechanisms driving pathogenicity in IDPRs, particularly in autosomal dominant genes, and propose strategies to enhance VEP accuracy by incorporating region-specific thresholds and IDPR-specific features. These findings provide a foundation for refining computational tools to better interpret the effects of genetic variants in disordered regions, with implications for understanding their role in human disease.

## Results and Discussion

### Pathogenic missense variants are depleted in disordered regions

To define intrinsically disordered regions, we used AlphaFold2 (AF2) pLDDT scores, which correlate inversely with structural order [16–18]. AF2 pLDDT is recognised as a robust predictor of disorder [17], demonstrating potential to surpass established disorder prediction models, as evidenced by benchmarking studies [19]. Residues were classified as disordered, ordered, or intermediate based on pLDDT and local context, as illustrated in Fig. 1A (see Methods). Importantly, our approach uses a conservative definition: only residues within long, low-confidence stretches (≥30 residues with average pLDDT <70) were classified as disordered. This threshold aims to capture regions likely to be truly unstructured under physiological conditions, such as extended flexible linkers or regulatory activation domains, rather than transiently mobile loops within structured domains. Shorter, low-confidence segments were instead assigned to an intermediate category to reduce false positives, particularly in cases where local flexibility does not imply global disorder. This strict classification likely underestimates the full extent of disorder in the proteome, but provides a high-confidence set of IDPRs for downstream analysis.

**Figure 1:**
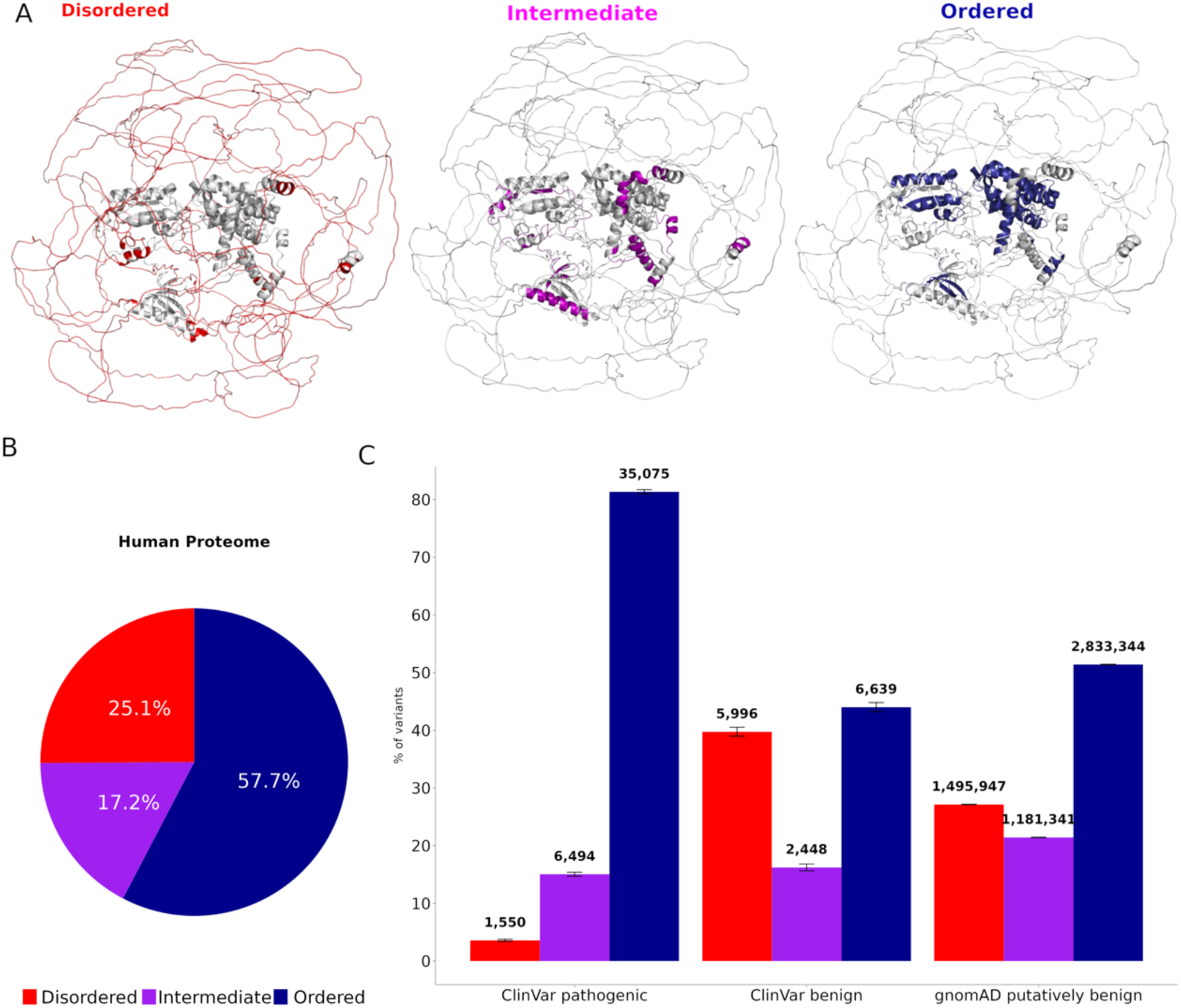
Rarity of pathogenic mutations in intrinsically disordered regions. (A) AF2-predicted structure of human Rho GTPase-activating protein 32 (UniProt ID: A7KAX9), with disordered, intermediate, and ordered regions depicted in red, purple, and blue, respectively. (B) Proportion of residues classified as disordered, intermediate, and ordered across the human proteome. (C) Distribution of missense variants across structural regions, expressed as percentages, with pathogenic and likely variants from ClinVar, benign variants from ClinVar, and putatively benign population variants from gnomAD.

Applying this approach to 20,281 canonical human proteins, we found that 25.1% of residues fall within disordered regions (Fig. 1B). In addition, 57.7% of residues are predicted to be ordered, while 17.2% are classified as intermediate. When considering disorder at the level of human proteins, most (57.7%) are predicted to have at least one disordered region, according to our definition, while 26.2% are highly disordered, having at least 30% of their residues in disordered regions (Figure S1). These observations are broadly consistent with previous findings [18,20], noting our strict disorder definition, as well as our more permissive intermediate classification.

We next examined the distribution of missense variants across structural regions. Variants were grouped into three categories: pathogenic and likely pathogenic variants from ClinVar [21], benign and likely benign variants from ClinVar, and putatively benign variants from gnomAD [22]. Pathogenic variants were strongly depleted in disordered regions, with only ∼3.6% occurring at disordered residues, compared to 15.1% in intermediate and 81.3% in ordered regions (Fig. 1C). In contrast, ClinVar benign variants were heavily enriched in disordered regions (39.8%). Finally, 27.2% of gnomAD putatively benign variants occurred in disordered regions, showing a slight enrichment over the occurrence of disorder in the human proteome. Overall, the vast majority of variants found in disordered regions are benign or putatively benign, while only a small minority are classified as pathogenic.

The strong enrichment of pathogenic missense variants in ordered regions and the corresponding enrichment of benign variants in disordered regions is broadly consistent with previous findings [23–25]. This distribution reflects fundamental differences in structural constraints between these regions. In a disordered region, a single amino acid substitution is unlikely to cause major functional disruption unless it affects a specific critical residue, such as one involved in PTMs or transient binding interactions. Conversely, in a structured domain, even a single missense variant can have a profound effect by destabilising the fold, altering interaction surfaces, or disrupting active sites. This difference in mutational tolerance aligns with the expectation that proteins rely on stable structural elements for core functions, whereas disordered regions often accommodate greater sequence variation without loss or change of function. Nevertheless, despite their depletion, we identified 1550 pathogenic missense variants in disordered regions, demonstrating that such variants can and do contribute to disease.

### Distinct mechanisms underlie pathogenic variants in disordered regions

Pathogenic missense variants in disordered regions must operate through distinct molecular mechanisms compared to those in ordered regions. In globular domains, such variants often cause disease by destabilising protein structure, leading to misfolding, or affecting the active site and ultimately a complete loss of function. In contrast, in disordered regions, pathogenic variants may affect protein behaviour by subtly altering regulatory interactions or disrupting specific binding motifs, such as short linear motifs or post-translational modification sites, without necessarily abolishing the protein’s overall function [26]. Given that current VEPs tend to perform best on loss-of-function (LOF) variants compared to gain-of-function (GOF) and dominant-negative (DN) variants [27], we can speculate that they might perform less well at the identification of pathogenic variants in disordered regions.

To explore this further, we first examined inheritance mode as a proxy for mechanism. Variants in autosomal recessive (AR) disorders nearly always act via LOF, while those in autosomal dominant (AD) disorders can involve LOF, GOF, or DN effects [28,29]. In Fig. 2A, we plot the proportion of variants in AD *vs* AR genes that occur in disordered, intermediate and ordered regions. Although the large majority of pathogenic variants are in ordered regions for both categories, it is interesting to note that AD genes show roughly double the proportion of pathogenic variants in disordered regions compared to AR genes (4.4% *vs* 2.2%, P = 3 x 10^-137^, Fisher’s exact test). Similarly, pathogenic variants in AD genes are also moderately enriched in intermediate regions (18.4% *vs* 11.9%, P = 1 x 10^-87^, Fisher’s exact test). This suggests that pathogenic variants in AR genes are more likely to act via a LOF mechanism involving structural disruption in an ordered region. In contrast, a much greater proportion of pathogenic variation in AD genes occurs in disordered regions, likely acting via more complex mechanisms related to the role of disordered regions in regulation, signalling, and interaction networks.

**Figure 2:**
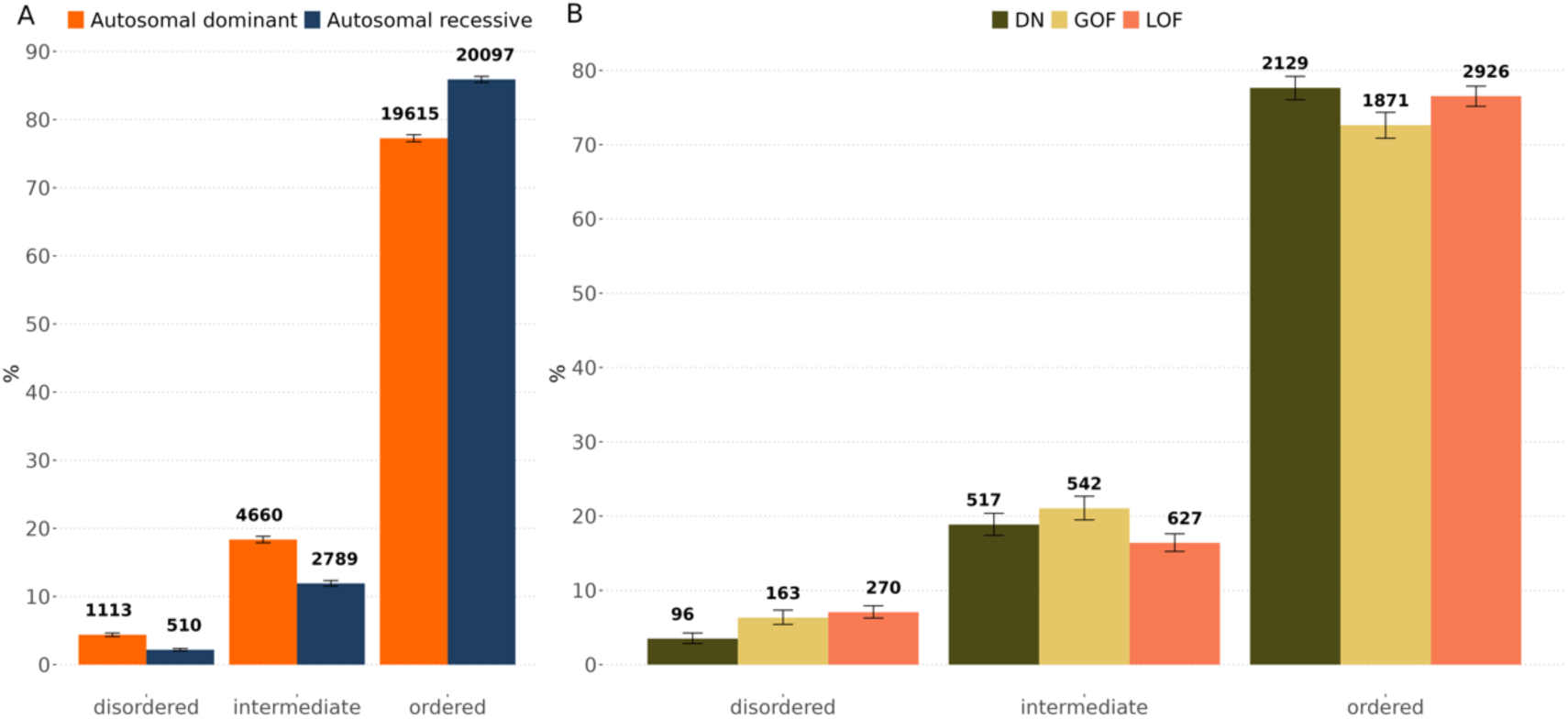
Distribution and counts of pathogenic missense variants by inheritance mode and molecular mechanism. (A) Bar plot illustrating the proportions and counts of pathogenic missense variants across disordered, intermediate, and ordered structural regions in autosomal dominant (AD) and autosomal recessive (AR) genes. AD genes exhibit approximately 2-fold and 1.5-fold higher proportions of pathogenic variants in disordered and intermediate regions, respectively, compared to AR genes. Counts of pathogenic variants per structural region and inheritance mode are indicated above each bar. (B) Bar plot showing the percentage and count of pathogenic missense variants associated with dominant negative (DN), gain-of-function (GOF), and loss-of-function (LOF) mechanisms within AD genes across structural regions. Counts are displayed above each bar, with error bars representing 95% binomial confidence intervals.

Next, we considered pathogenic variants from AD genes, and grouped them by molecular mechanism (LOF, GOF and DN) used previously performed literature-based classification [27] (Fig. 2B). Surprisingly, LOF variants showed the highest enrichment in disordered regions, in marked contrast with our initial expectation based on the results of the AR *vs* AD variants, and our naïve expectation of how pathogenic variants in disordered regions would be likely to act. Interestingly, however, LOF variants showed the lowest representation in intermediate regions. On closer consideration, we noted that many of the disordered LOF variants in AD genes occurred in transcription factors, which have a very well-known association with haploinsufficiency [30]. Given that many transcription factors contain large, disordered regions, often playing important roles in transactivation [31,32], this suggests that disruption of these functions is a common mechanism for pathogenicity of variants in disordered regions.

GOF variants were also enriched in disorder compared to AD variants in general, with 6.3% occurring at predicted disordered regions. Interestingly, GOF variants also showed the strongest enrichment at intermediate regions, and the lowest representation in ordered regions. This is consistent with the previous observation that GOF variants were enriched in regions with lower pLDDT values [27]. This pattern is consistent with the flexible, context-dependent nature of disordered and intermediate regions, which often mediate transient, regulated interactions. GOF variants in these regions may create new or stronger interactions, disrupt phase separation, interfere with post-translational modification sites, or perturb short linear motifs involved in dynamic regulation. These changes can enhance or misdirect protein activity without necessarily compromising stability or folding, making disordered and intermediate regions plausible sites for such effects.

Finally, only 2.2% of DN variants occurred at disordered regions, much lower than for the other AD mechanisms. This is likely due to the strong association of the DN effect with protein oligomerisation, and the tendency for DN variants to be at or close to intersubunit interfaces [27], which will generally tend to be ordered. Thus, the DN effect is not likely to be a common mechanism for pathogenicity in disordered regions, although there are plausible mechanisms, *e.g.* via competitive DN effects that do not require oligomersiation [28].

### VEP performance varies across structural regions

Next, we investigated how VEPs perform in distinguishing pathogenic from putatively benign missense variants across disordered, intermediate and ordered regions. We used 33 different VEPs, with strict coverage filters, so that all VEPs had predictions for all variants tested (see Methods). Although this limits the number of VEPs we can include, it means that our results will not be influenced any coverage biases of individual methods, given that many VEPs do not make predictions available for all genes, or even for all variants within the genes they cover [13].

We grouped into three categories using a recently introduced classification scheme [33,34], based on their potential risk of circularity that arises when evaluating VEPs on variants potentially related to those used in training [35]. *Clinical-trained* VEPs are supervised models trained directly on human variants with known clinical labels, such as pathogenic and benign annotations. These inherently have the highest risk of circularity. *Population-tuned* VEPs are not trained on clinical labels but have been optimised or calibrated using *human population* data, typically through allele frequency-based scaling or tuning. These tend to have a much lower, but non-zero susceptibility to circularity. Finally, *population-free* VEPs have not been trained or tuned on any human variant data and are therefore immune from circularity concerns. This group includes unsupervised methods, protein language models, and models based on evolutionary conservation from sequence alignments.

In Figure 3, we show the AUROC values for disordered, intermediate and ordered regions across all predictors. In Fig. 3A, we group population-free and population-tuned VEPs together (noting that most are population-free and only AlphaMissense [36], UNEECON [37] and LIST-S2 [38] are population-tuned). Fig. 3B shows the clinical-trained VEPs, which comprise the majority of methods in our analysis. Interestingly, nearly all VEPs show the highest AUROC values in disordered regions and the lowest values in ordered regions. The only exception to this is MPC [39], which shows slightly worse performance in disordered than ordered regions, though we note that its performance across all regions remains poor compared to other VEPs.

**Figure 3:**
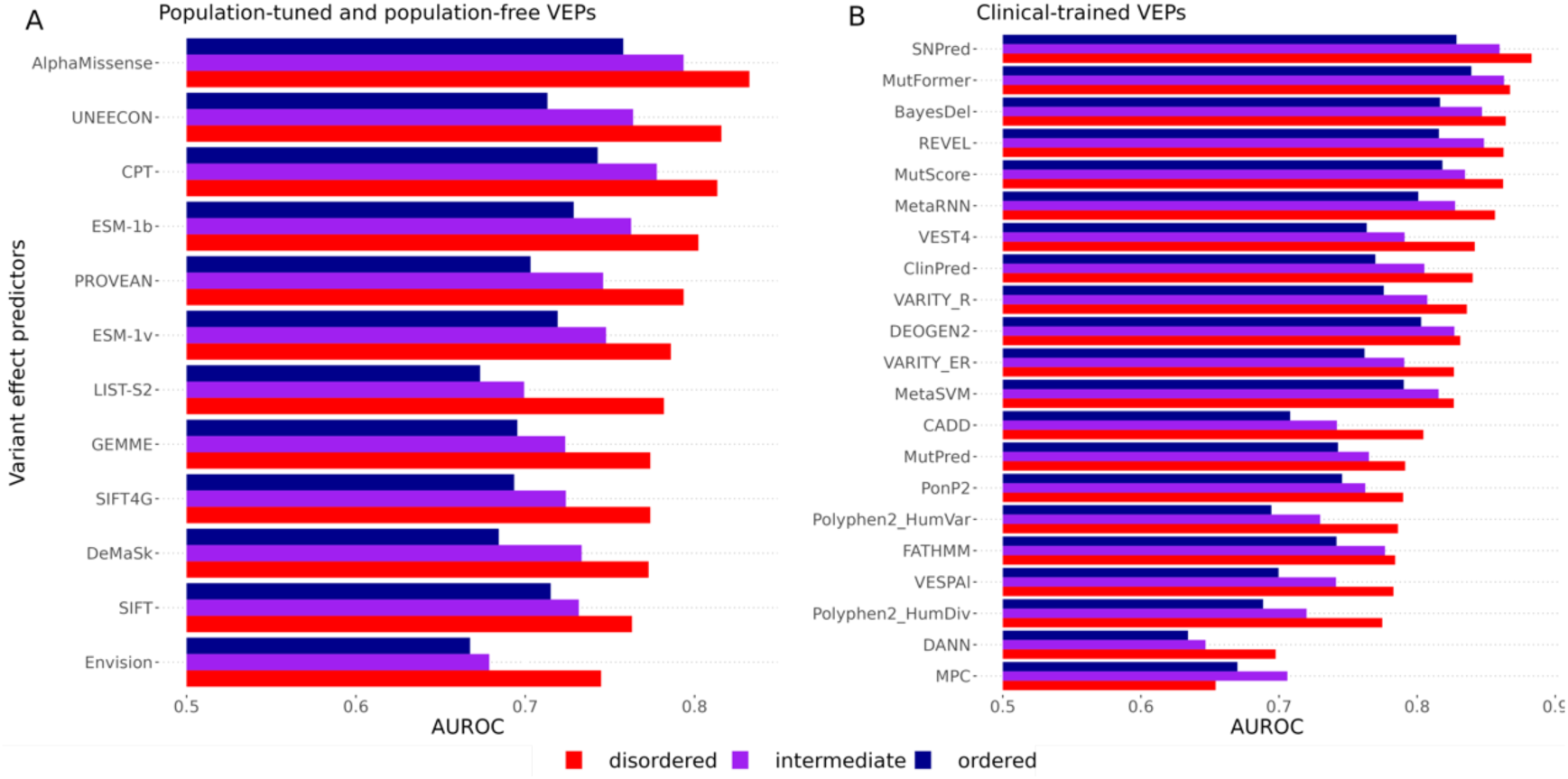
Performance evaluation of VEPs across structural regions. Performance of VEPs assessed by area under the receiver operating characteristic curve (AUROC) across disordered, intermediate, and ordered protein regions. The x-axis represents AUROC values starting at 0.5, and the y-axis lists individual VEPs. Population-free and population-tuned VEPs are shown together in (A), while clinical-trained VEPs are shown in (B).

When comparing different VEPs, there is little evidence that any methods perform particularly well in specific regions. For example, AlphaMissense and CPT [40] show the highest AUROC of any population-tuned or population-free models across disordered, intermediate and ordered regions. Similarly, SNPred [41] and MutFormer [42] show higher AUROC values than any clinical-trained models across all three regions. Thus, methods that perform well on ordered regions also tend to perform well on disordered regions, and there does not seem to be any reason to recommend a particular VEP for ordered *vs* disordered regions. Notably, VEPs based on protein language models (*e.g.* ESM-1b [43] and ESM-1v [44]) appear to show very similar trends across regions as those based purely on sequence alignments (*e.g.* GEMME [45]), suggesting that neither approach has any clear advantages in ordered *vs* disordered regions. This is somewhat surprising, as protein language models have been proposed to capture contextual sequence signals that might help in disordered regions where structural information is lacking [43]. However, our findings suggest that in practice, these models do not currently offer enhanced predictive power in IDPRs.

### VEPs show low sensitivity for pathogenic variants in disordered regions

The observation that VEPs show superior performance in disordered regions, as measured by AUROC, is consistent with our previous study, where we found that the presence of large disordered regions was highly predictive of AUROC, and that excluding disordered regions tended to reduce AUROC values [15]. There we noted that this phenomenon was driven not by the properties of pathogenic variants, but by the high density of putatively benign variants in disordered regions, which tend to be weakly conserved and thus are more easily classified as benign by most VEPs.

To investigate this phenomenon further, we calculated the sensitivity and specificity at the “optimal threshold” point of ROC curve for each VEP (see Methods). The sensitivity represents the true positive rate: the fraction of pathogenic variants that are correctly classified as pathogenic using this threshold. The specificity represents the true negative rate: the fraction of putatively benign variants correctly classified as non-pathogenic.

In Fig. 4A, we plot sensitivity *vs* specificity for all VEPs in disordered, intermediate and ordered regions. Strikingly, across nearly all predictors, sensitivity for pathogenic variants in disordered regions was significantly lower compared to ordered and intermediate regions. This low sensitivity indicates that, despite high overall AUROC scores driven by accurate identification of putatively benign variants, VEPs frequently misclassify pathogenic variants within disordered sequences. At the same time, specificities are clearly much higher in disordered regions than ordered regions. That is, while VEPs are less likely to correctly classify variants as pathogenic in disordered regions, they are more likely to misclassify benign variants.

**Figure 4:**
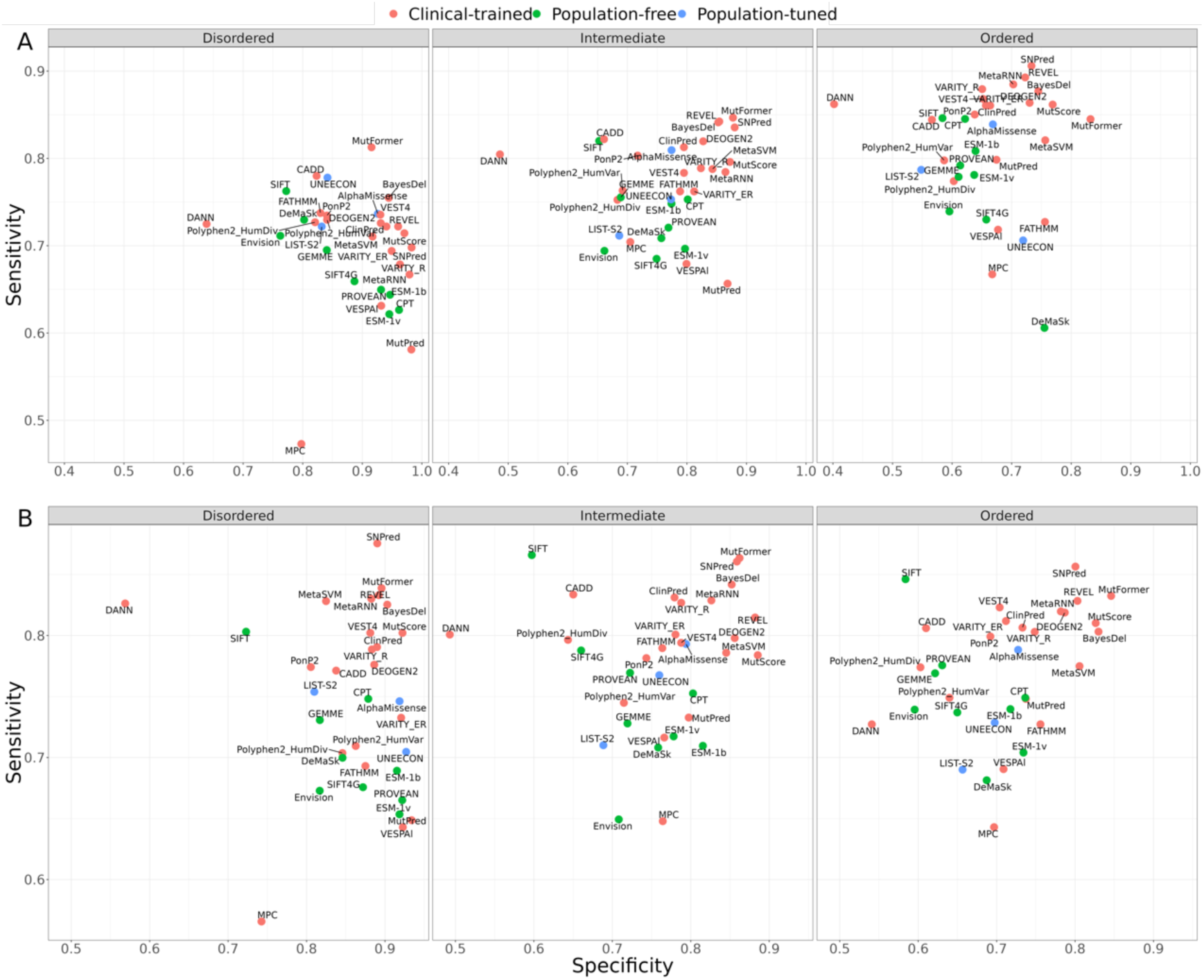
Assessment of VEP performance using global and region-specific optimal thresholds. Performance of VEPs evaluated by sensitivity on the x-axis and specificity on the y-axis. (A) Sensitivity *vs* specificity in disordered, intermediate and ordered regions calculated using a global optimal threshold, across all structural regions. (B) Sensitivity *vs* specificity calculated using a region-specific optimal threshold.

We observe that clinical-trained VEPs tend to have higher sensitivities than population-free VEPs, although this is almost certainly driven by the circularity issue discussed earlier. Interestingly, however, we also note a strong inverse correlation between sensitivity and specificity for population-free VEPs within disordered regions. While some degree of trade-off between these metrics is expected, one plausible explanation is that population-free predictors rely predominantly on evolutionary conservation or biophysical signals that are inherently weaker in disordered regions. As a result, adjusting their decision threshold to capture more pathogenic variants (thus boosting sensitivity) inadvertently increases misclassification of benign variants, reducing specificity. Conversely, a more stringent threshold that accurately dismisses most benign variants sharply lowers sensitivity for the sparse, but functionally impactful, pathogenic variants in IDPRs.

Given that the low sensitivity of VEPs in disordered regions means that they are likely to miss true pathogenic variants, we wondered whether the use of region-specific thresholds could be beneficial. Thus, we calculated the optimal thresholds for each VEP from the ROC curves in the same manner as before, but considering only disordered, intermediate or ordered variants (Table S1). Essentially, this results in lower thresholds for classifying variants as pathogenic in disordered regions, and higher thresholds in ordered regions (assuming a VEP where the score positively correlates with likelihood of pathogenicity). For example, for CPT, the top-performing population-free VEP, we calculate a global optimal threshold of 0.35, compared to a disordered-specific threshold of 0.24 and ordered-specific threshold of 0.44. Note that these are not meant to be thresholds for making clinical classifications; far stricter thresholds would be required for this [46]. Instead, these represent the optimal thresholds for discriminating between pathogenic and putatively benign in our dataset, and therefore suggest useful thresholds for consideration in variant prioritisation.

In Fig. 4B, we plot sensitivity *vs* specificity using these region-specific thresholds, while Figure S3 shows the difference in sensitivity and specificity using region-specific vs global thresholds. The use of region-specific thresholds results in very similar sensitivities of VEPs across all three regions, generally increasing sensitivity in disordered regions and decreasing it in ordered regions. Interestingly, while specificity is also affected, the impact appears to be relatively smaller than on sensitivity in disordered regions This suggests that the sensitivity gain in disordered regions is likely worth the relatively small impact on specificity. In contrast, the loss of sensitivity in ordered regions is not compensated for by a large specificity increase. Thus, we suggest that these region-specific thresholds are likely useful for disordered and possibly intermediate regions, but they may be of less benefit for prioritising variants in ordered regions.

Given that that evolutionary conservation plays a key role in most VEPs, we wondered whether the reduced sensitivity of VEPs could be related to reduced conservation of pathogenic variants in disordered regions. We address this in Figure S2, comparing the residue-level conservation of the sites of pathogenic and putatively benign variants in disordered, intermediate and ordered regions. Unsurprisingly, pathogenic variants occur at far more conserved positions than putatively benign variants. Interestingly, however, both pathogenic and putatively benign variants in disordered regions are less conserved residues than those in ordered regions. Thus, the weaker conservation of pathogenic variants in disordered regions means they are less likely to be correctly classified, thus reducing sensitivity, while the weaker conservation of putatively benign variants means that they are more likely to be correctly classified, thus increasing specificity.

### VEPs show discordant predictions in disordered regions

Given the observed variability in VEP performance across structural contexts, we next investigated the consistency of their predictions for individual variants. Specifically, we asked whether VEPs tend to agree on which variants are pathogenic in intrinsically disordered regions (IDPRs), compared to intermediate and ordered regions.

To quantify agreement, we calculated the average pairwise concordance between VEPs using Cohen’s kappa statistic. This metric measures inter-rater reliability while correcting for agreement expected by chance. A kappa value of 1 indicates perfect agreement, 0 corresponds to agreement no better than random, and negative values reflect systematic disagreement. This allows us to assess not just whether two predictors classify variants similarly, but whether they do so more consistently than would be expected by chance alone. We binarised VEP outputs using each method’s global optimal threshold (see Methods), and calculated mean pairwise kappa values within and between VEP groups defined by training strategy—clinical-trained, population-tuned, and population-free.

As shown in Figure 5, overall agreement was lowest in disordered regions, particularly among population-free VEPs. These models, which rely primarily on evolutionary conservation and sequence-derived features, appear to disagree more frequently in regions lacking strong structural or conservation signals. This suggests that the sparse and subtle functional constraints characteristic of IDPRs lead to reduced model convergence and greater uncertainty in prediction.

**Figure 5:**
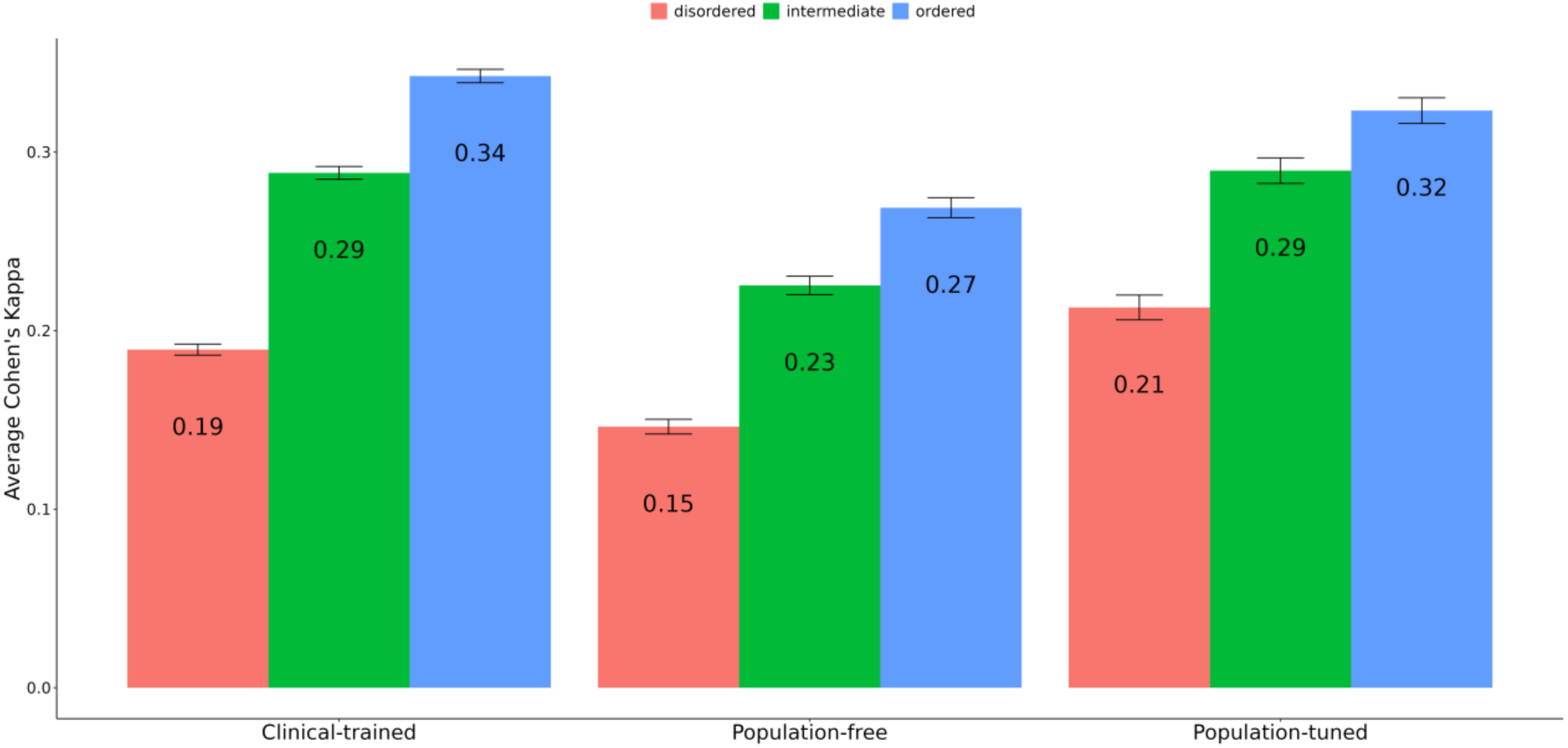
Average classification agreement among VEP groups across structural regions. Average Cohen’s kappa values representing classification agreement among VEP groups across disordered, intermediate, and ordered protein structural regions. The x-axis lists VEP groups categorised by training methodology, and the y-axis indicates the mean Cohen’s kappa for each group within each structural region. Error bars denote the standard error of the mean.

In contrast, clinical-trained and population-tuned VEPs showed somewhat higher agreement in disordered regions. This likely reflect shared biases arising from human variant data used during model development or calibration, which can lead to convergence due to circularity rather than true predictive power. Nonetheless, even within these groups, agreement was lower in disordered regions than in ordered regions.

By comparison, all VEP groups showed higher concordance in intermediate and ordered regions, consistent with the stronger evolutionary and structural constraints in these contexts. Here, models are more likely to converge on consistent classifications, likely driven by clearer functional signals.

Together, these findings suggest that disordered regions not only pose challenges for sensitivity but also reduce agreement between VEPs, particularly those without access to human variant data. This highlights the need for specialised strategies—both in training and interpretation—to improve consistency and reliability in these structurally flexible regions.

## Conclusion

IDPRs are crucial mediators of diverse regulatory functions, yet their inherent structural flexibility poses significant challenges for accurate interpretation of genetic variants associated with human disease. Our systematic analysis highlights fundamental limitations in current VEPs, which demonstrate notably reduced sensitivity for pathogenic variants within these disordered contexts. Although VEPs consistently achieve higher AUROC scores in IDPRs, this largely reflects their proficiency at identifying benign variants—driven by weaker evolutionary constraints—and not their ability to pinpoint subtle yet functionally critical pathogenic alterations. Consequently, relying solely on global thresholds risks overlooking clinically relevant variants that operate through mechanisms unique to disordered regions.

We demonstrate that pathogenic variants in IDPRs predominantly act through distinct molecular mechanisms compared to those in structured domains, particularly involving nuanced alterations in regulatory interactions or transcriptional activity. These mechanisms are often inadequately captured by current predictors, especially those relying heavily on evolutionary conservation signals or structured-domain assumptions. Our findings strongly advocate for incorporating region-specific thresholds into VEPs to enhance their sensitivity and accuracy for disordered regions without substantially compromising specificity.

Furthermore, the notable discordance among predictors in classifying IDPR variants underscores the importance of refining computational methodologies and training strategies tailored specifically for disordered protein regions. Future VEP development could potentially achieve improved performance and reliability by incorporating region-specific features such as transient binding motifs, post-translational modification sites, and context-dependent structural ensembles. Recent large-scale simulations of human IDRs demonstrate that pathogenic variants are enriched in regions with low conformational entropy, providing a new structural dimension for interpreting variant effects in disordered regions [47]. In parallel, recent proteome-wide analyses have highlighted that pathogenic variants in disordered regions more often act via direct disruption of function rather than through loss of stability, and that such variants are frequently misclassified by current predictors, reflecting both the distinct mechanisms at play and the limitations of models trained predominantly on folded proteins [48].

Overall, this work provides critical insights into the complexities of interpreting genetic variation within intrinsically disordered regions. It establishes a foundational understanding for future development of more sophisticated computational tools, ultimately enhancing the accuracy of genetic variant interpretation in clinical settings and broadening our comprehension of disease mechanisms rooted in protein disorder.

## Methods

### Structural classification of human residues

Every residue in the human proteome, considering the primary UniProt isoform of each protein-coding gene was given a structural classification of ordered, intermediate or disordered. To do this, we utilised the pLDDT derived from AF2 [16]. pLDDT inversely correlates very well with the flexibility of protein structures such that AF2 assigns low pLDDT scores for regions that are highly flexible and lack fixed 3D structures such as IDPRs and linkers between ordered regions [17,18]. A residue is deemed to belong to a disordered region if its pLDDT is less than 50 and it is part of a contiguous stretch of at least 30 residues with an average pLDDT value less than 70. In contrast, a residue is classified as ordered if its pLDDT is at least 70. Finally, residues falling outside these two conditions were classified as intermediate.

### Missense variant dataset

Our dataset of missense variants was derived in essentially the same way as in our previous study [15], with pathogenic and likely pathogenic variants taken from ClinVar (August 2022) [21], and putatively benign population variants from gnomAD v2.1 [22]. Using gnomAD as a source of ‘putatively benign’ variants is preferable to clinically classified benign variants because it reduces circularity: many VEPs incorporate allele frequency information, which is often used to label clinical benign variants [49], creating a risk of inflated performance due to overlap between training and evaluation data. In contrast, using mostly rare gnomAD variants better reflects the real-world challenge of distinguishing rare benign from rare pathogenic variants while offering a larger, less biased negative class [33,50]. However, we also separately considered benign variants from ClinVar in Figure 1. Importantly, we also removed all missense variants from collagen proteins, by excluding those 213 human protein-coding genes containing collagen-helices, as defined by Pfam (PF01391) [51]. It has been demonstrated previously that presence of collagen-helix containing proteins skew analyses of intrinsic disorder, since they are fibrous proteins but are consistently predicted to be disordered [52]. In our initial analysis, we found that nearly half of pathogenic missense variants in “disordered” regions according to our definition were from collagen proteins. As we do not consider collagens to be intrinsically disordered proteins, we excluded them from our study, and we strongly suggest that people take this into consideration in future studies of disease variants in in disordered regions, as they have a high potential to cause confounding.

### Assessing VEP performance and agreement

We started with the set of VEPs tested in a recent benchmarking study (considering those used in the original preprint) [33]. We only retained those methods with scores available for at least 75% of missense variants present in our pathogenic and putatively benign datasets. To ensure consistency of comparisons, we only retained variants with scores shared across all VEPs.

For each VEP, we assessed its performance at distinguishing between pathogenic and putatively benign variants by calculating the AUROC value across different structural regions. We also calculated “optimal thresholds” for distinguishing between pathogenic and putatively benign variants, either on a global basis, or considering specific structural regions. To do this, we applied the Youden J-statistic [53] to the ROC curve and selected the threshold that maximised this value [53]. Using these thresholds, we could then assess whether each VEP predicted each variant to be pathogenic or benign in a binary manner. From this, we could calculate other metrics, including sensitivity and specificity. We also used this to calculate Cohen’s kappa, to assess the level of agreement in classification between each pair of VEPs.

## Data availability

Associated datasets are available at https://doi.org/10.6084/m9.figshare.c.7747895.v1 and the pipeline code is shared at https://github.com/drsamibioinfo/VEPS_IN_DISORDER/

## Acknowledgments

We thank Benjamin Livesey and Mihaly Badonyi for helpful comments on the manuscript. This project was supported by funding to JAM from the European Research Council (ERC) under the European Union’s Horizon 2020 research and innovation programme (grant agreement No. 101001169) and by the Medical Research Council (MRC) Human Genetics Unit core grant (MC_UU_00035/9). This work has made use of the resources provided by the Edinburgh Compute and Data Facility (ECDF) (http://www.ecdf.ed.ac.uk/).

## Supplementary Material

**Figure S1:**
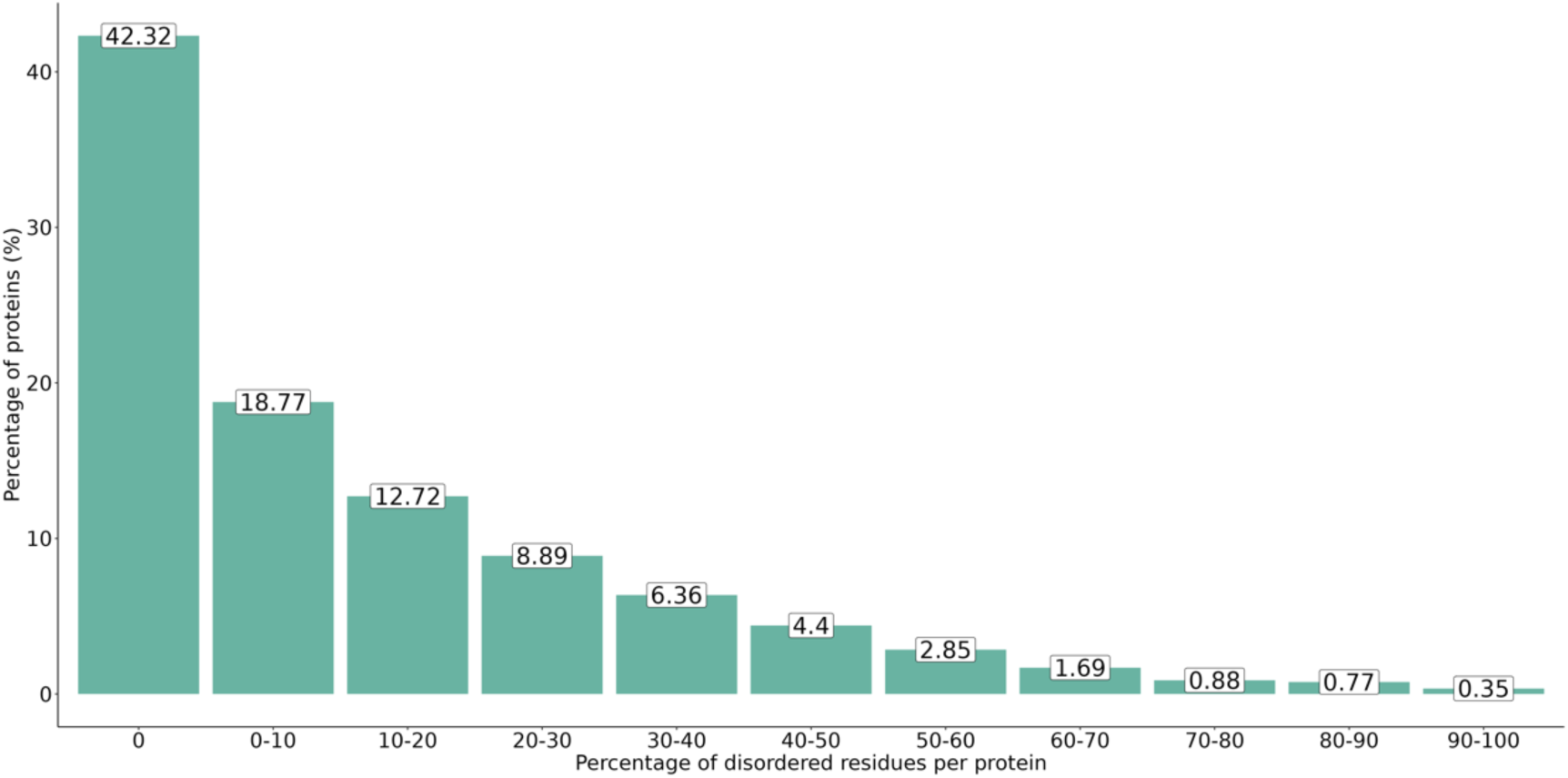
Distribution of human proteins by percentage of intrinsic disorder. Proportion of human proteins across varying levels of IDPRs. The x-axis represents the percentage of IDPRs, grouped into bins, and the y-axis shows the percentage of human proteins within each bin. Bins are left-inclusive and right-exclusive (*e.g*., 10.5% falls in the “10–20” bin), while the “0%” bin includes only fully ordered proteins.

**Figure S2:**
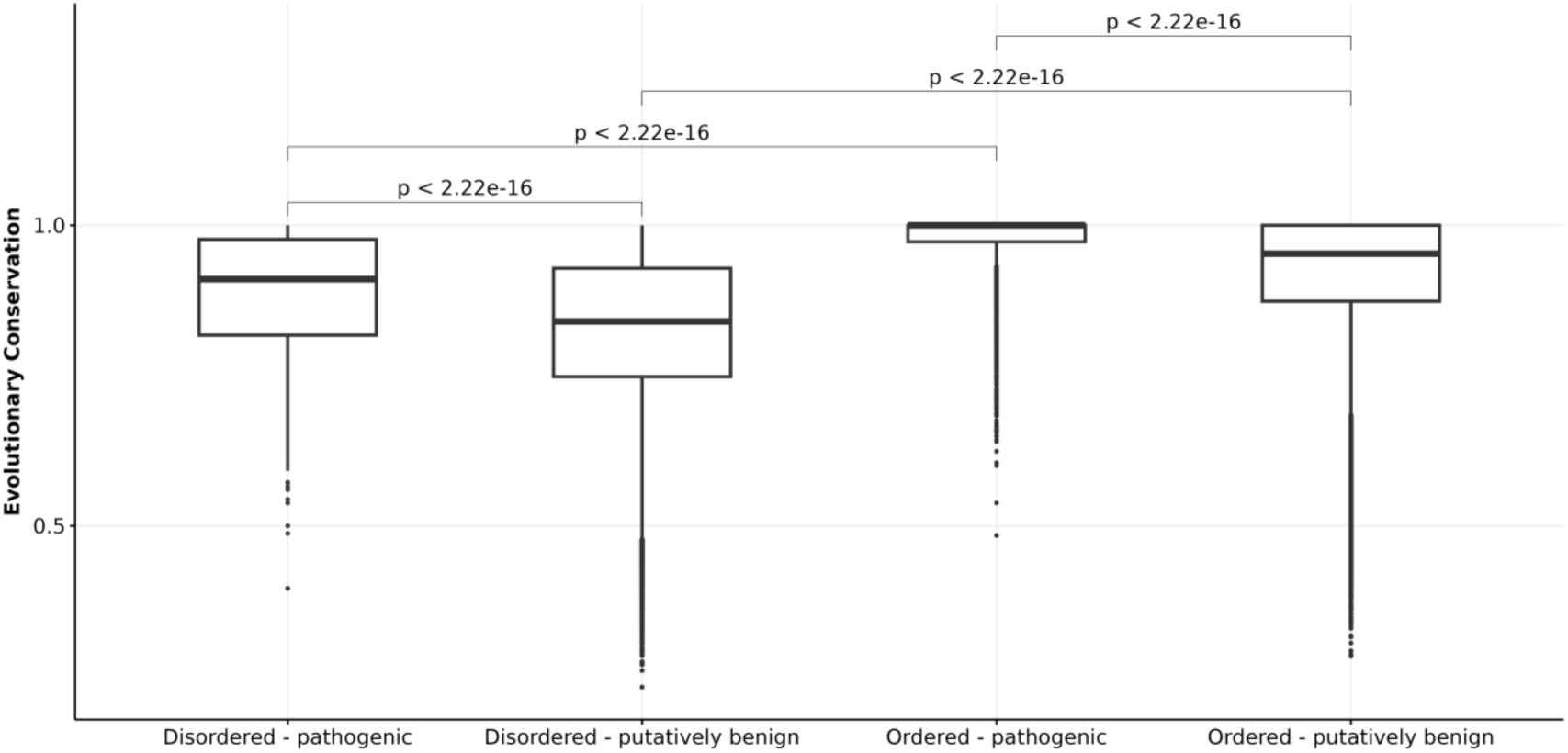
Evolutionary conservation of variants in disordered and ordered regions: Comparison of evolutionary conservation between pathogenic and putatively benign variants in disordered and ordered protein regions. The x-axis denotes the functional consequence of variants (pathogenic or putatively benign) within each structural region. The y-axis represents evolutionary conservation, calculated as the complement of the relative residue substitution rate (1 - substitution rate) from multiple sequence alignments of closely related homologues using Aminode, with higher values indicating greater residue conservation. P-values, calculated using the Wilcoxon rank-sum test, are displayed for comparisons between groups as indicated.

**Figure S3:**
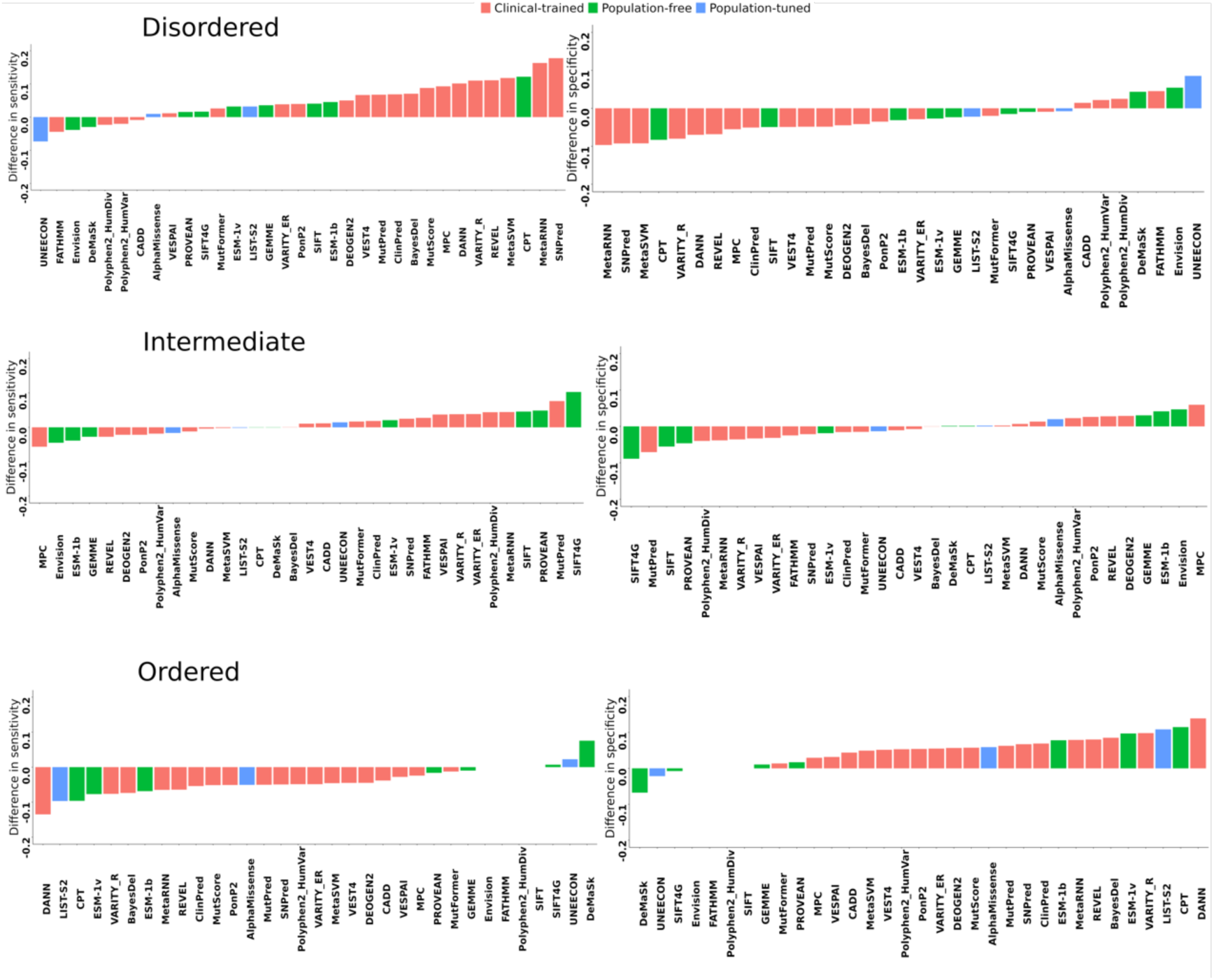
Difference in sensitivity and specificity of VEPs using region-specific *vs* global optimal thresholds. A sensitivity or specificity difference > 0 indicates that employing the region-specific optimal threshold results in higher sensitivity or specificity the global optimal threshold.

**Table S1:** Global and region-specific optimal thresholds across all VEPs.

| VEP | Gobal threshold | Disordered threshold | Intermediate threshold | Ordered threshold |
| --- | --- | --- | --- | --- |
| AlphaMissense | 0.5204 | 0.486 | 0.5701 | 0.6468 |
| BayesDel | 0.153387 | 0.0833305 | 0.152938 | 0.232318 |
| CADD | 24.15 | 24.3666667 | 24.05 | 24.56667 |
| ClinPred | 0.94754 | 0.88315082 | 0.938846111 | 0.97119 |
| CPT | 0.352497 | 0.24362566 | 0.353933498 | 0.439558 |
| DANN | 0.991105 | 0.98662852 | 0.991432234 | 0.995396 |
| DeMaSk | -0.256 | -0.2802 | -0.2568 | -0.219 |
| DEOGEN2 | 0.439909 | 0.338799 | 0.491653 | 0.519703 |
| Envision | 0.868355 | 0.854226 | 0.858515 | 0.868355 |
| ESM-1b | -9.15 | -7.889 | -9.835 | -10.041 |
| ESM-1v | -7.861874 | -6.9753757 | -7.56275148 | -8.959092 |
| FATHMM | -1.135 | -1.78 | -0.94 | -1.14 |
| GEMME | -1.991826 | -1.8200497 | -2.16361472 | -2.04416 |
| LIST-S2 | 0.89581 | 0.885811 | 0.89651 | 0.925307 |
| MetaRNN | 0.786569 | 0.46689788 | 0.73198164 | 0.852458 |
| MetaSVM | -0.06 | -0.5516 | -0.0472 | 0.1494 |
| MPC | 0.732077 | 0.61975996 | 0.855313714 | 0.784322 |
| MutFormer | 0.666125 | 0.5343537 | 0.5865506 | 0.7227 |
| MutPred | 0.641 | 0.497 | 0.571 | 0.681 |
| MutScore | 0.6555 | 0.466 | 0.683 | 0.739 |
| Polyphen2_HumDiv | 0.984 | 0.992 | 0.965 | 0.984 |
| Polyphen2_HumVar | 0.821 | 0.885 | 0.868 | 0.905 |
| PonP2 | 0.553 | 0.495 | 0.589 | 0.62 |
| PROVEAN | -3.305 | -3.15 | -2.92 | -3.43 |
| REVEL | 0.500667 | 0.338 | 0.5535 | 0.61 |
| SIFT | 0.0101 | 0.02 | 0.02 | 0.01 |
| SIFT4G | 0.0125 | 0.015 | 0.03 | 0.013 |
| SNPred | 0.88215 | 0.5205498 | 0.851998276 | 0.932433 |
| UNEECON | 0.243275 | 0.363272 | 0.231323 | 0.226351 |
| VARITY_ER | 0.49784 | 0.40675068 | 0.4414843 | 0.580362 |
| VARITY_R | 0.577377 | 0.33660173 | 0.50747865 | 0.723914 |
| VESPAI | 0.47849 | 0.46934984 | 0.460013334 | 0.493212 |
| VEST4 | 0.704 | 0.61 | 0.696 | 0.752 |

## Notes

https://doi.org/10.6084/m9.figshare.c.7747895.v1

### Competing Interest Statement

The authors have declared no competing interest.

## References

1. Das RK, Pappu RV. Conformations of intrinsically disordered proteins are influenced by linear sequence distributions of oppositely charged residues. Proc Natl Acad Sci. 2013;110: 13392–13397. doi:10.1073/pnas.1304749110

2. Lermyte F. Roles, Characteristics, and Analysis of Intrinsically Disordered Proteins: A Minireview. Life. 2020;10: 320. doi:10.3390/life10120320

3. Pajkos M, Dosztányi Z. Functions of intrinsically disordered proteins through evolutionary lenses. Progress in Molecular Biology and Translational Science. Elsevier; 2021. pp. 45–74. doi:10.1016/bs.pmbts.2021.06.017

4. Kurgan L, Hu G, Wang K, Ghadermarzi S, Zhao B, Malhis N, et al. Tutorial: a guide for the selection of fast and accurate computational tools for the prediction of intrinsic disorder in proteins. Nat Protoc. 2023;18: 3157–3172. doi:10.1038/s41596-023-00876-x

5. Lazar T, Tantos A, Tompa P, Schad E. Intrinsic protein disorder uncouples affinity from binding specificity. Protein Sci. 2022;31: e4455. doi:10.1002/pro.4455

6. Lee TY, Huang HD, Hung JH, Huang HY, Yang YS, Wang TH. dbPTM: an information repository of protein post-translational modification. Nucleic Acids Res. 2006;34. doi:10.1093/nar/gkj083

7. Uversky V. p53 Proteoforms and Intrinsic Disorder: An Illustration of the Protein Structure–Function Continuum Concept. Int J Mol Sci. 2016;17: 1874. doi:10.3390/ijms17111874

8. Uversky VN. Protein intrinsic disorder and structure-function continuum. Progress in Molecular Biology and Translational Science. Elsevier; 2019. pp. 1–17. doi:10.1016/bs.pmbts.2019.05.003

9. Uversky VN, Oldfield CJ, Dunker AK. Intrinsically Disordered Proteins in Human Diseases: Introducing the D ^2^ Concept. Annu Rev Biophys. 2008;37: 215–246. doi:10.1146/annurev.biophys.37.032807.125924

10. Dunker AK, Lawson JD, Brown CJ, Williams RM, Romero P, Oh JS, et al. Intrinsically disordered protein. J Mol Graph Model. 2001;19: 26–59.

11. Uversky VN. What does it mean to be natively unfolded?: Natively unfolded proteins. Eur J Biochem. 2002;269: 2–12. doi:10.1046/j.0014-2956.2001.02649.x

12. Vovk A, Zilman A. Effects of Sequence Composition, Patterning and Hydrodynamics on the Conformation and Dynamics of Intrinsically Disordered Proteins. Int J Mol Sci. 2023;24: 1444. doi:10.3390/ijms24021444

13. Livesey BJ, Badonyi M, Dias M, Frazer J, Kumar S, Lindorff-Larsen K, et al. Guidelines for releasing a variant effect predictor. arXiv; 2024. doi:10.48550/arXiv.2404.10807

14. Vacic V, Markwick PRL, Oldfield CJ, Zhao X, Haynes C, Uversky VN, et al. Disease-Associated Mutations Disrupt Functionally Important Regions of Intrinsic Protein Disorder. Ofran Y, editor. PLoS Comput Biol. 2012;8: e1002709. doi:10.1371/journal.pcbi.1002709

15. Fawzy M, Marsh JA. Understanding the heterogeneous performance of variant effect predictors across human protein-coding genes. Sci Rep. 2024;14: 26114. doi:10.1038/s41598-024-76202-6

16. Jumper J, Evans R, Pritzel A, Green T, Figurnov M, Ronneberger O, et al. Highly accurate protein structure prediction with AlphaFold. Nature. 2021;596. doi:10.1038/s41586-021-03819-2

17. Tunyasuvunakool K, Adler J, Wu Z, Green T, Zielinski M, Žídek A, et al. Highly accurate protein structure prediction for the human proteome. Nature. 2021;596. doi:10.1038/s41586-021-03828-1

18. Alderson TR, Pritišanac I, Kolarić Đ, Moses AM, Forman-Kay JD. Systematic identification of conditionally folded intrinsically disordered regions by AlphaFold2. Proc Natl Acad Sci U S A. 2023;120: e2304302120. doi:10.1073/pnas.2304302120

19. Wilson CJ, Choy W-Y, Karttunen M. AlphaFold2: A Role for Disordered Protein/Region Prediction? Int J Mol Sci. 2022;23: 4591. doi:10.3390/ijms23094591

20. Pentony MM, Jones DT. Modularity of intrinsic disorder in the human proteome. Proteins Struct Funct Bioinforma. 2010;78: 212–221. doi:10.1002/prot.22504

21. Landrum MJ, Lee JM, Riley GR, Jang W, Rubinstein WS, Church DM, et al. ClinVar: Public archive of relationships among sequence variation and human phenotype. Nucleic Acids Res. 2014;42. doi:10.1093/nar/gkt1113

22. Karczewski KJ, Francioli LC, Tiao G, Cummings BB, Alföldi J, Wang Q, et al. The mutational constraint spectrum quantified from variation in 141,456 humans. Nature. 2020;581. doi:10.1038/s41586-020-2308-7

23. Feng M, Wei X, Zheng X, Liu L, Lin L, Xia M, et al. Decoding Missense Variants by Incorporating Phase Separation via Machine Learning. Nat Commun. 2024;15: 8279. doi:10.1038/s41467-024-52580-3

24. Lee KE, Pulido JS, Da Palma MM, Procopio R, Hufnagel RB, Reynolds M. A Comprehensive Report of Intrinsically Disordered Regions in Inherited Retinal Diseases. Genes. 2023;14: 1601. doi:10.3390/genes14081601

25. Wong ETC, So V, Guron M, Kuechler ER, Malhis N, Bui JM, et al. Protein–Protein Interactions Mediated by Intrinsically Disordered Protein Regions Are Enriched in Missense Mutations. Biomolecules. 2020;10: 1097. doi:10.3390/biom10081097

26. Iakoucheva LM, Brown CJ, Lawson JD, Obradović Z, Dunker AK. Intrinsic disorder in cell-signaling and cancer-associated proteins. J Mol Biol. 2002;323: 573–584.

27. Gerasimavicius L, Livesey BJ, Marsh JA. Loss-of-function, gain-of-function and dominant-negative mutations have profoundly different effects on protein structure. Nat Commun. 2022;13. doi:10.1038/s41467-022-31686-6

28. Backwell L, Marsh JA. Diverse Molecular Mechanisms Underlying Pathogenic Protein Mutations: Beyond the Loss-of-Function Paradigm. Annu Rev Genomics Hum Genet. 2022;23: 475–498. doi:10.1146/annurev-genom-111221-103208

29. Badonyi M, Marsh JA. Proteome-scale prediction of molecular mechanisms underlying dominant genetic diseases. PloS One. 2024;19: e0307312. doi:10.1371/journal.pone.0307312

30. Seidman JG, Seidman C. Transcription factor haploinsufficiency: when half a loaf is not enough. J Clin Invest. 2002;109: 451–455. doi:10.1172/JCI15043

31. Boija A, Klein IA, Sabari BR, Dall’Agnese A, Coffey EL, Zamudio AV, et al. Transcription Factors Activate Genes through the Phase-Separation Capacity of Their Activation Domains. Cell. 2018;175: 1842–1855.e16. doi:10.1016/j.cell.2018.10.042

32. Wright PE, Dyson HJ. Intrinsically disordered proteins in cellular signalling and regulation. Nat Rev Mol Cell Biol. 2015;16: 18–29.

33. Livesey BJ, Marsh JA. Variant effect predictor correlation with functional assays is reflective of clinical classification performance. bioRxiv; 2024. p. 2024.05.12.593741. doi:10.1101/2024.05.12.593741

34. Pathak AK, Bora N, Badonyi M, Livesey BJ, Consortium S, Ngeow J, et al. Pervasive ancestry bias in variant effect predictors. bioRxiv; 2025. p. 2024.05.20.594987. doi:10.1101/2024.05.20.594987

35. Grimm DG, Azencott CA, Aicheler F, Gieraths U, Macarthur DG, Samocha KE, et al. The evaluation of tools used to predict the impact of missense variants is hindered by two types of circularity. Hum Mutat. 2015;36. doi:10.1002/humu.22768

36. Cheng J, Novati G, Pan J, Bycroft C, Žemgulytė A, Applebaum T, et al. Accurate proteome-wide missense variant effect prediction with AlphaMissense. Science. 2023;381: eadg7492. doi:10.1126/science.adg7492

37. Huang Y-F. Unified inference of missense variant effects and gene constraints in the human genome. PLOS Genet. 2020;16: e1008922. doi:10.1371/journal.pgen.1008922

38. Malhis N, Jacobson M, Jones SJM, Gsponer J. LIST-S2: taxonomy based sorting of deleterious missense mutations across species. Nucleic Acids Res. 2020;48: W154– W161. doi:10.1093/nar/gkaa288

39. Samocha KE, Kosmicki JA, Karczewski KJ, O’Donnell-Luria AH, Pierce-Hoffman E, MacArthur DG, et al. Regional missense constraint improves variant deleteriousness prediction. bioRxiv; 2017. p. 148353. doi:10.1101/148353

40. Jagota M, Ye C, Albors C, Rastogi R, Koehl A, Ioannidis N, et al. Cross-protein transfer learning substantially improves disease variant prediction. Genome Biol. 2023;24: 182. doi:10.1186/s13059-023-03024-6

41. Molotkov I, Koboldt DC, Artomov M. SNPred outperforms other ensemble-based SNV pathogenicity predictors and elucidates the challenges of using ClinVar for evaluation of variant classification quality. medRxiv; 2023. p. 2023.09.07.23295192. doi:10.1101/2023.09.07.23295192

42. Jiang TT, Fang L, Wang K. Deciphering “the language of nature”: A transformer-based language model for deleterious mutations in proteins. The Innovation. 2023;4: 100487. doi:10.1016/j.xinn.2023.100487

43. Brandes N, Goldman G, Wang CH, Ye CJ, Ntranos V. Genome-wide prediction of disease variant effects with a deep protein language model. Nat Genet. 2023;55: 1512–1522. doi:10.1038/s41588-023-01465-0

44. Meier J, Rao R, Verkuil R, Liu J, Sercu T, Rives A. Language models enable zero-shot prediction of the effects of mutations on protein function. Advances in Neural Information Processing Systems. Curran Associates, Inc.; 2021. pp. 29287–29303. Available: https://proceedings.neurips.cc/paper/2021/hash/f51338d736f95dd42427296047067694-Abstract.html

45. Laine E, Karami Y, Carbone A. GEMME: A Simple and Fast Global Epistatic Model Predicting Mutational Effects. Mol Biol Evol. 2019;36: 2604–2619. doi:10.1093/molbev/msz179

46. Pejaver V, Byrne AB, Feng B-J, Pagel KA, Mooney SD, Karchin R, et al. Calibration of computational tools for missense variant pathogenicity classification and ClinGen recommendations for PP3/BP4 criteria. Am J Hum Genet. 2022;109: 2163. doi:10.1016/j.ajhg.2022.10.013

47. Tesei G, Trolle AI, Jonsson N, Betz J, Knudsen FE, Pesce F, et al. Conformational ensembles of the human intrinsically disordered proteome. Nature. 2024;626: 897– 904. doi:10.1038/s41586-023-07004-5

48. Cagiada M, Jonsson N, Lindorff-Larsen K. Decoding molecular mechanisms for loss of function variants in the human proteome. bioRxiv; 2024. p. 2024.05.21.595203. doi:10.1101/2024.05.21.595203

49. Richards S, Aziz N, Bale S, Bick D, Das S, Gastier-Foster J, et al. Standards and Guidelines for the Interpretation of Sequence Variants: A Joint Consensus Recommendation of the American College of Medical Genetics and Genomics and the Association for Molecular Pathology. Genet Med Off J Am Coll Med Genet. 2015;17: 405–424. doi:10.1038/gim.2015.30

50. Wu Y, Li R, Sun S, Weile J, Roth FP. Improved pathogenicity prediction for rare human missense variants. Am J Hum Genet. 2021;108: 1891–1906. doi:10.1016/j.ajhg.2021.08.012

51. Paysan-Lafosse T, Andreeva A, Blum M, Chuguransky SR, Grego T, Pinto BL, et al. The Pfam protein families database: embracing AI/ML. Nucleic Acids Res. 2025;53: D523– D534. doi:10.1093/nar/gkae997

52. Smithers B, Oates ME, Tompa P, Gough J. Three reasons protein disorder analysis makes more sense in the light of collagen. Protein Sci. 2016;25: 1030–1036. doi:10.1002/pro.2913

53. Youden WJ. Index for rating diagnostic tests. Cancer. 1950;3: 32–35. doi:10.1002/1097-0142(1950)3:1<32::AID-CNCR2820030106>3.0.CO;2-3

